# Long range projections of oxytocin neurons in the marmoset brain

**DOI:** 10.1101/2024.01.02.573953

**Authors:** Arthur Lefevre, Jazlynn Meza, Cory T. Miller

## Abstract

The neurohormone oxytocin (OT) has become a major target for the development of novel therapeutic strategies to treat psychiatric disorders such as autism spectrum disorder because of its integral role in governing many facets of mammalian social behavior. Whereas extensive work in rodents has produced much of our knowledge of OT, we lack basic information about its neurobiology in primates making it difficult to interpret the limited effects that OT manipulations have had in human patients. In fact, previous studies have revealed only limited OT fibers in primate brains. Here, we investigated the OT connectome in marmoset using immunohistochemistry, and mapped OT fibers throughout the brains of adult male and female marmoset monkeys. We found extensive OT projections reaching limbic and cortical areas that are involved in the regulation of social behaviors, such as the amygdala, the medial prefrontal cortex and the basal ganglia. The pattern of OT fibers observed in marmosets is notably similar to the OT connectomes described in rodents. Our findings here contrast with previous results by demonstrating a broad distribution of OT throughout the marmoset brain. Given the prevalence of this neurohormone in the primate brain, methods developed in rodents to manipulate endogenous OT are likely to be applicable in marmosets.

## Introduction

Oxytocin (OT) is a neurohormone involved in the regulation of many physiological functions in the brain that has a profound effect on a number of behaviors, most notably facets of sociality [1]. Because of the putative role that OT plays in a number of social disorders in humans [2], significant efforts have been made to develop novel therapeutic strategies that modulate its regulation. And yet despite our growing understanding of the neurobiological circuits underlying the actions of OT in rodents [1,3,4], most clinical trials using OT in human patients have failed to produce conclusive effects [5,6]. While the reason for this discrepancy is not clear, there are at least two explanations that may account for the pattern of results. First, some functions of OT system may be different between rodents and humans owing to their evolutionary divergence of primates 100 million years ago. Second, the effects of precise OT manipulations done in rodents with tools such as opto- and chemogenetics or local intracerebral OT injection are not recapitulated by the methods commonly used in human and non-human primates, such as intranasal administration, blood measurements or genetic association studies [7–13]. While settling these methodological issues will take time, a critical first step towards addressing this gap of knowledge is to compare the anatomy of the OT system between species and determine shared and divergent properties. Unfortunately, to this date, the distributions of OT fibers and OT receptor in the primate brain have been only partially studied [14–20], thereby precluding our ability to infer the functions of OT neuronal pathways identified in rodents [21–27] to human and nonhuman primates.

The limited number of studies seeking to identify OT fibers in the primate brain have yielded somewhat different and conflicting results. Extra hypothalamic OT fibers have not been consistently reported and found mostly around the hypothalamus and brainstem in marmosets [16] and macaques [14], and only one study reported OT axons in cortical areas [28]. The available data thus show a very different picture to the patterns reported in rodents [29,30]. Here we performed a whole brain quantitative analysis of OT fibers in male and female common marmoset monkeys (Callithrix jacchus) to bridge the gap in our knowledge of this keystone neurohormone in a primate species by validating and applying a new OT antibody during immunohistochemical processing. Marmosets have emerged as a powerful neurobiological model of primate brain function, particularly for facets of social brain functions [31]. This New World primate pair-bonds, cooperatively cares for their young, and engage in a suite of prosocial behaviors including food-sharing, imitation and social gaze [32,33]. Moreover, these monkeys are highly voluble, frequently engaging in bouts of vocal exchanges [34]. The neurobiological basis of vocal communication in marmosets is the most thoroughly studied of all social brain functions in this primate [35–39], and this function may be modulated by OT. Thus, the many similar social characteristics shared between marmosets and humans make this species particularly intriguing to examine the prevalence of OT fibers in the brain of a nonhuman primate.

## Methods

### Animals

The brains from four adult common marmosets (2 males and 2 females, aged 1y11mo, 2y4mo, 3y1mo and 7y1mo) were used in this study. Animals had to be euthanized for various reasons (2 for wasting syndrome, 1 for implant loss and 1 for AAV virus investigation). Before euthanasia, all animals were group housed, and experiments were performed in the Cortical Systems and Behavior Laboratory at University of California San Diego (UCSD). All experiments were approved by the UCSD Institutional Animal Care and Use Committee.

### Perfusion and tissue processing

Animals were anesthetized with Ketamine and then euthanized with pentobarbital sodium. Phosphate Buffer Saline (PBS) was perfused transcardially until all the blood was flushed out of the body, followed by 200mL of 4% paraformaldehyde (PFA). The brain was quickly removed and postfixed overnight in 4% PFA, before being stored at 4°C in PBS with 0.025% sodium azide until the tissue was cut using a vibratome in 50µm thick coronal slices.

### Immunohistochemistry

Immunohistochemistry was performed on free-floating sections. For each brain, 1 in 6 slice (1 slice every 250µm) was processed. Following 3 washes with PBS, slices were preincubated for 2 hours in 1% Triton and 2% NGS on PBS, incubated overnight at room temperature on a plate shaker with 1:5000 oxytocin rabbit antibody (T-4084, Peninsula lab) and/or 1:1000 vasopressin guinea pig antibody (403004, SySy). The day after, slices were washed 3 x 10 minutes with PBS, incubated in the dark for 2 hours with fluorescent secondary antibody Cy3 (111 165 003, Jackson IR) and/or alexa A488 (706 545 148, Jackson IR), washed again with PBS then mounted on Superfrost glass slides and coverslipped using Vectashield mounting medium with DAPI (H-1200-10, Vector laboratories).

For control experiments, 1µg of synthetic OT (O4375, Sigma Aldrich) or AVP (V0377, Sigma Aldrich) were added to the well during incubation with primary antibody. It should be noted that this synthetic OT is derived from human OT and thus has a different amino acid at the 8 position (New World monkeys have a proline instead of leucine [40]), however, this did not seem influence our results nor antibody specificity. We also co-stained AVP and OT to ensure the specificity of the OT antibody.

### Imaging and analysis

Slides were imaged using a slide scanner epifluorescent microscope (Axioscan 7, Zeiss) and digitalized automatically. OT fibers were manually counted and attributed to a brain region according to the Paxinos atlas [41]. A total of 3628 individual fibers were counted across the 4 brains. Fibers within the hypothalamus were not quantified but qualitatively described because of their high density due to the OT cell bodies being in proximity. Results are shown as percentage obtained by dividing the number of OT fibers found in one brain area by the total number of extra hypothalamic OT fibers. We only integrated an area in our analysis if it was found to contain OT fibers in all 4 analyzed brains or 2 of the same sex. Due to lower tissue quality and processing, the brainstem was not analyzed by counting fibers but by qualitative assessment (see Fig 4).

## Results

To provide a whole brain characterization of OT in the marmoset monkey, we quantified the prevalence of these fibers from the brain slices of four adult marmosets (2male/2female). We first verified the specificity of the OT antibody by performing competition experiments on adjacent slices containing the paraventricular nucleus of the hypothalamus (PVN). During incubation with the primary OT antibody, we added synthetic OT (1µg), synthetic vasopressin (AVP) (1µg) or PBS. We found that artificial OT prevented the OT antibody from labelling OT neurons in the PVN (Fig 1a) but that this was not the case when adding artificial AVP (Fig 1b) or PBS (Fig 1c). This validates the specificity of the OT antibody used in this study with marmosets. Furthermore, we also co stained OT and AVP in the same slice (Fig 1d) and found no co-labelling, confirming that the OT antibody does not cross react with AVP in marmosets.

**Figure 1.**
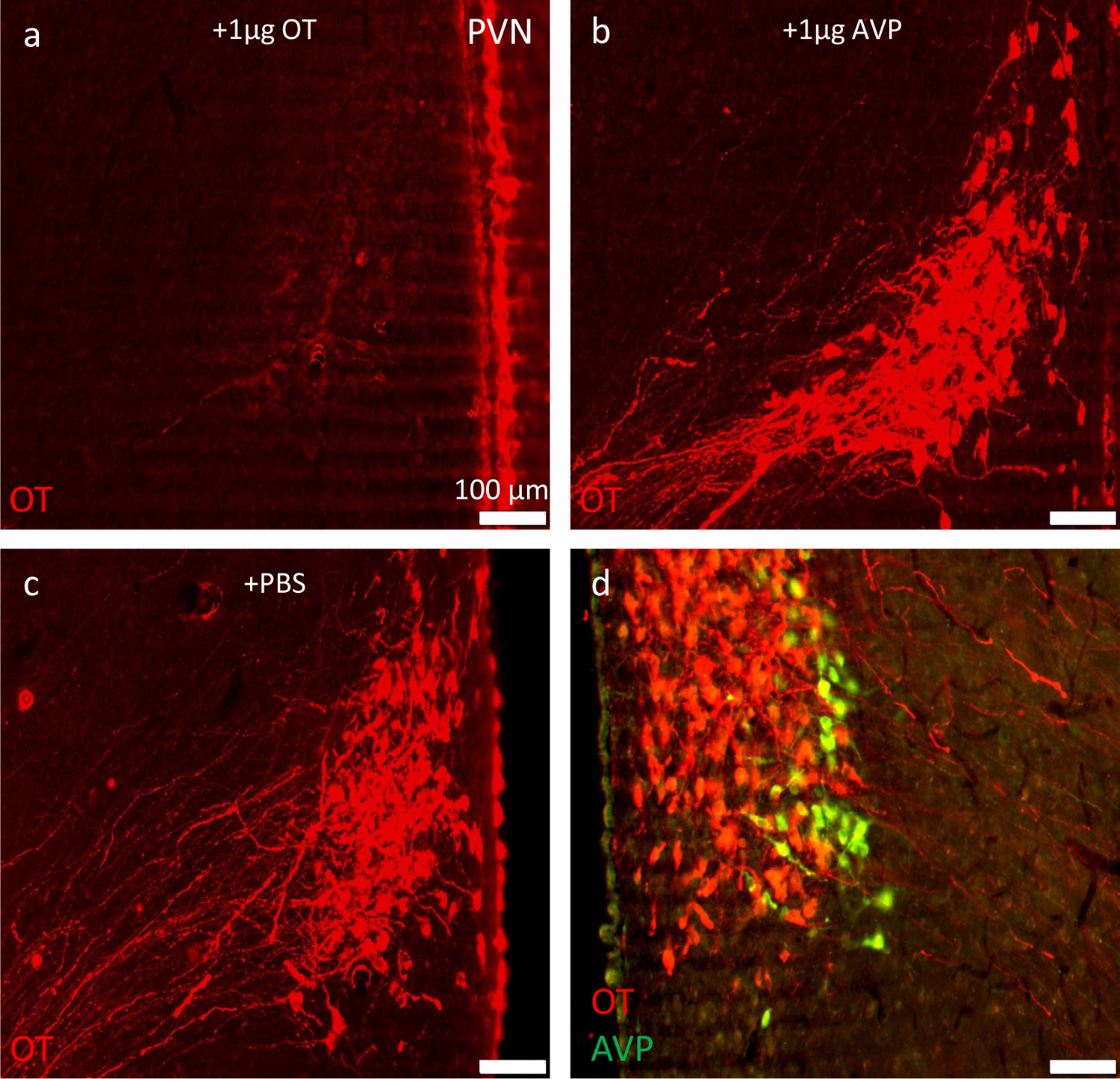
OT staining of a slice containing PVN incubated with 1µg of artificial OT **(a)**, 1µg of artificial AVP **(b)** or saline **(c)**. All images were taken with the same acquisition parameters (light intensity, gain, exposure, color brightness and balance). **(d)** Co staining for OT (in red) and AVP (in green) in the same slice.

Our initial analysis sought to determine the organization of OT neurons in the marmoset hypothalamus. Consistent with previously described [16] pattern and to the Paxinos atlas [41], we identified the PVN from interaural +10.5mm, a little anterior to the anterior commissure, to interaural +7.8mm (Fig 2). The supraoptic nucleus (SON) was likewise located between interaural +11.0mm and +8.5mm. The retrochiasmatic part of the SON was evident between interaural +9.0mm and +7.8mm (Fig 2). As expected [16], we observed high densities of OT neurons in both the PVN and SON (Fig 2), while a small number of OT positive cells were also evident in the Stria Terminalis close to the ventricle junction (lateral and third) around interaural +8.6mm. Finally, few OT cells were seen in between PVN and SON, between interaural +9.5mm and +8.0mm, but as opposed to rodents [21,42,43] no accessory nucleus was observed in marmosets (Fig 2).

**Figure 2.**
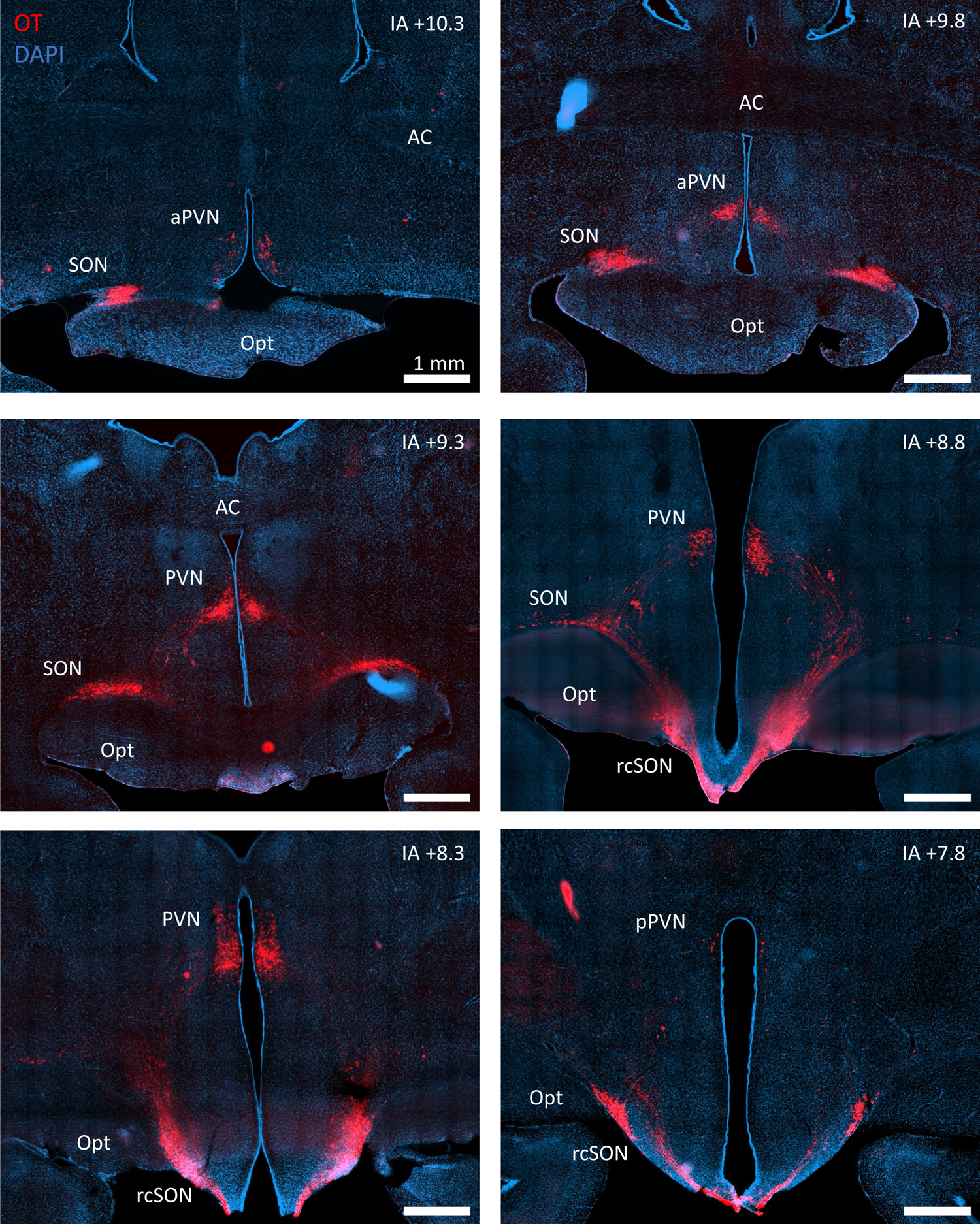
Fluorescent photomicrographs of coronal hypothalamic slices containing OT neurons (in red). IA: interaural level, AC: anterior commissure, Opt: optic tract.

Because of the paucity of data on the distribution of OT fibers in the primate brain, we next systematically quantified OT fibers through the brains of all four marmoset subjects. This analysis revealed OT fibers are evident in considerably more areas of the primate brain than had previously been reported [14,16]. First, we observed high innervation (not quantified) of all hypothalamic areas, including the preoptic area (median, medial and lateral), the anterior, lateral, ventro- and dorso-medial, and posterior hypothalamus as well as the mammillary body and zona incerta. At the sub cortical level, we found a dense plexus in the endopiriform nucleus that extended to the claustrum (Fig 3), as well as medium innervation to the nucleus accumbens (Fig 3). In addition, we found several regions close to the hypothalamus with an elevated OT fibers count such as the amygdala (in both central and basolateral parts), the bed nucleus of stria terminalis, the horizontal diagonal band of Broca and Meynert nucleus (Fig 3). Interestingly we also found several cortical areas that contained

**Figure 3.**
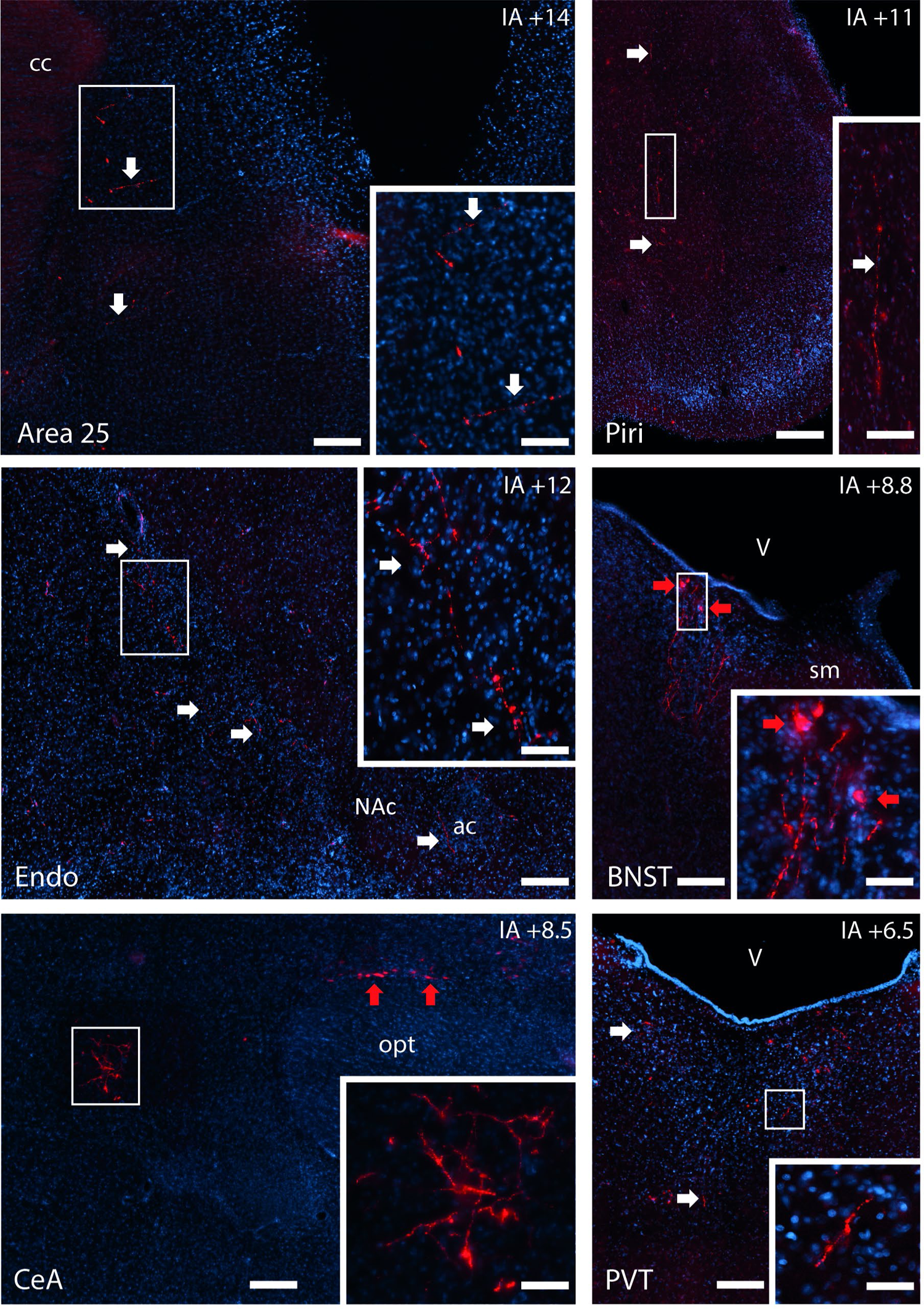
Photomicrographs of coronal slices containing OT fibers (in red). Fibers are indicated by white arrows, cell bodies by red arrows (found in BNST and SON). Scale bar = 300µm (100µm for insets). IA: interaural level in mm, ac: anterior commissure, BNST: bed nucleus of the stria terminalis, cc: corpus callosum, CeA: central amygdala, Endo: endopiriform nucleus, Opt: optic tract, Piri: piriform cortex, PVT: paraventricular nucleus of the thalamus, sm: stria medullaris, V: Ventricle.

OT fibers. These included mostly median areas like the anterior cingulate cortex (such as area 24, 25 and 32) and ventrolateral areas in the orbital and temporal cortices (such as area 13, 14, temporal areas, entorhinal and insula). Finally, in the posterior half of the brain, light but consistent OT innervation was evident in the brain stem, mostly directed to the periaqueductal gray with sparser fibers to other nuclei such as the inferior and superior colliculus, the nucleus of tractus solitary and the locus coeruleus. We did not observe differences between the male and female brains nor between left and right hemispheres (see fig 4a). A comprehensive list of brain regions found innervated by OT fibers can be found in Fig 4a and a representative schema in Fig 4b.

**Figure 4.**
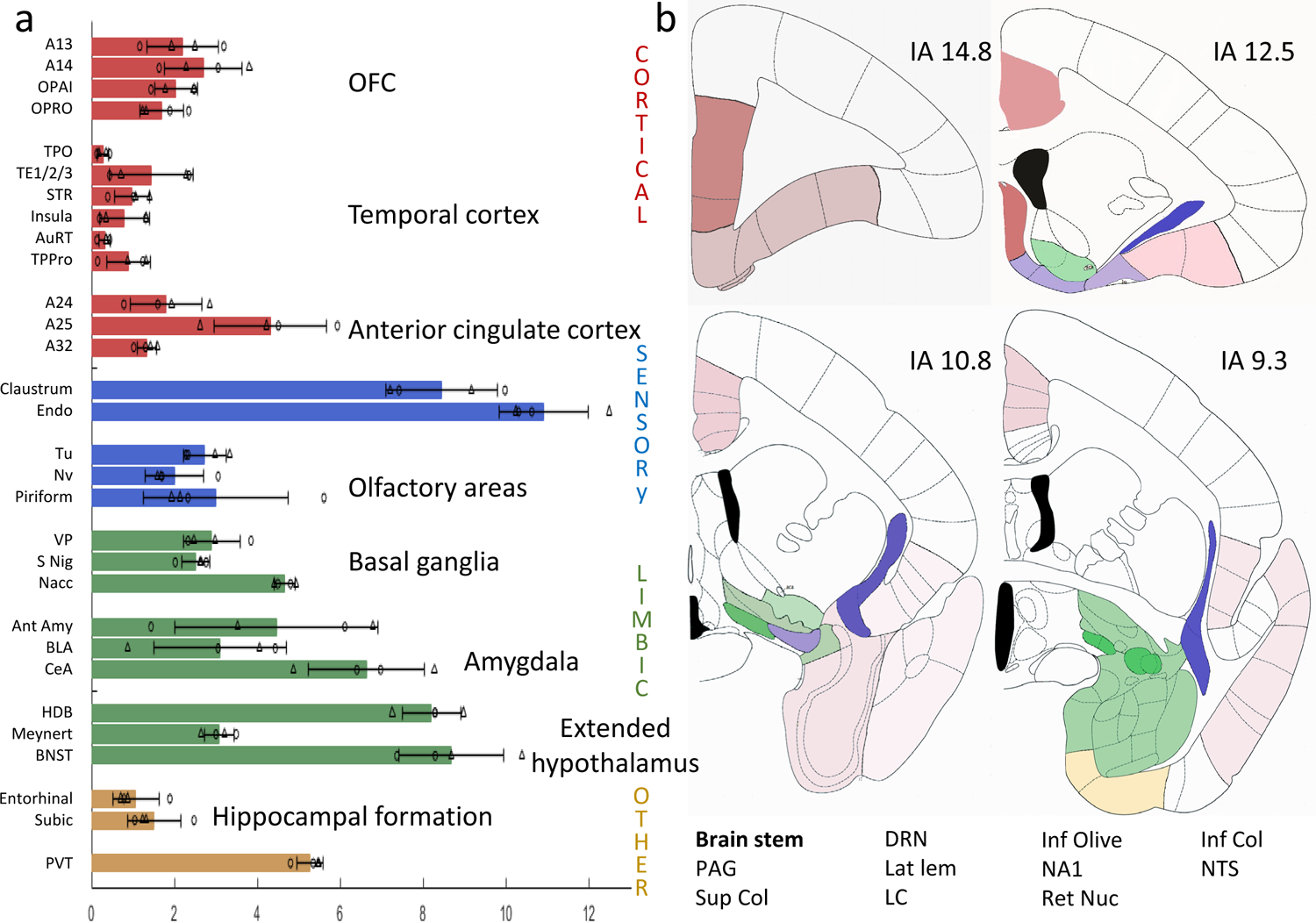
**(a)** Graph indicating individual (triangles and circles represent males and females, respectively) and averaged level of OT innervation per region, expressed as percent of the total. Error bars represent standard deviation. Lower right: list of brainstem regions containing OT fibers (not quantified). **(b)** Schematics of OT fibers distribution on four representative coronal sections (IA: Inter Aural level in mm), colors indicate the type of region (same as in (a), cortical in red, sensory in blue, limbic in green and others in yellow) and color intensity indicates fiber density. Abbreviations: Ant Amy: anterior amygdala, AuRT: auditory rostrotemporal cortex, BLA: basolateral amygdala, BNST: bed nucleus of the stria terminalis, CeA: central amygdala, DRN: dorsal raphe nucleus, Endo: endopiriform nucleus, HDB: horizontal limb of the diagonal band, Inf Oli: inferior olive, Lat lem: lateral lemniscus, LC: locus coeruleus, NA1: noradrenergic group 1, NAcc: nucleus accumbens, NTS: nucleus of the solitary tract, Nv: navilar nucleus, OPAI: orbital periallocortex, OPro: orbital proisocortex, PAG: periaqueductal gray, Piri: piriform cortex, PVT: paraventricular nucleus of the thalamus, Ret nuc: reticular nucleus, S Nig: substancia nigra, STR: superior temporal rostral area, Subic: subiculum, Sup/Inf col: superior/inferior colliculus, TE1/2/3: temporal areas 1/2/3, TPO: temporal par occ assoc, TPPro: Temporopolar proisocortex, Tu: olfactory tubercle, VP: Ventral pallidum. Adapted from the Paxinos atlas [41].

## Discussion

OT is synthesized mostly by neurons in two nuclei of the hypothalamus, the paraventricular nucleus (PVN) located close to the third ventricle and the supraoptic nucleus (SON) located above the optic tract [44,45], an organization that is highly conserved across vertebrate taxa [40], including humans [15]. In the present study, we investigated and mapped OT fibers throughout the male and female marmoset brain and observed OT fibers in numerous brain regions, including multiple cortical areas. We observed numerous extra hypothalamic OT fibers, especially in the anterior half of the brain, most of which were found close to the midline (ACC, BNST, PVT, PAG and brainstem), ventral (OFC, Olf, HDB, basal ganglia, Meynert nucleus, amydgala) or ventro temporal (claustrum, hippocampal formation, temporal cortex). In the posterior half of the brain, we only found OT fibers in the PAG and the brainstem, with a very similar distribution to what was previously found in macaques [14] and rodents [30], albeit the lower quality of data from the brainstem prevented us to quantify the fibers. Although no significant differences were observed between males and females, subtle differences may not have been evident because of the small number of marmoset brains analyzed, and the nature of the analysis performed here. It should however be noted that sex and hemispheric differences in rodents are only evident when comparing virgin and pregnant/lactating animals [21,22,46], and both of our female subjects were nulliparous. We also cannot rule out false negatives from our antibody which could lead to underestimation or lack of detection of some OT fibers. Other techniques, such as AAV anterograde tracing, are needed to confirm these results. The widespread prevalence of OT fibers throughout the brain notably contrasts with previous studies in marmosets [16] and macaques [14], perhaps due to newer and more sensitive antibodies or more systematic investigation, and opens the door to more direct comparisons with rodents.

The pattern of OT projections observed in marmosets here is consistent with analogous studies in rodents [21,25,29,30], with the notable exception of lateral septum. While this structure is well innervated with OT fibers in rats [47], none were detected in marmoset lateral septum. The consistent similarities in OT fibers across taxa are of critical importance because we can now hypothesize that these shared pathways have similar functions in rodents and primates. For instance, OT release in the amygdala is known to have anxiolytic effects [21] but also to be important for social recognition [48]. In the nucleus accumbens, OT is involved in social reward [49] and regulates pair bonding in voles [4]. It would thus be very interesting to manipulate OT in these areas in marmosets, as these highly social monkeys also display pair bonding. Furthermore, recent studies have demonstrated that in rodents, OT in medial prefrontal cortical areas is critical for social motivation and sociability, notably in the median prefrontal cortex [50,51]. This finding is potentially significant because the human mPFC (area 24, 25, 32) has been found to be a hub for neuropeptidergic transmission [52]. Finally, it should be noted that some of the most innervated areas in both rodents and marmosets such as the PVT and the endopiriform nucleus have been scantly studied. The largely conserved neurobiology of OT between rodents and marmosets provides a roadmap in which to more precisely interrogate the functional role of these pathways in a primate to improve therapeutic treatments for the myriad disorders affected by OT given that intranasal OT administration has yielded such poor results.

Many of the brain areas innervated by OT in marmosets are involved in vocal communication, such as auditory and temporal cortices, ACC areas 32 and 24 and amygdala. As OT is important for social behaviors [1] and marmosets are known to be highly social and voluble [34], the occurrence of these fibers in these neural substrates is perhaps not surprising but demonstrates an important link between this neurohormone and multiple social brain structures in primates. This pattern is also consistent with the theory that OT and oxytocin receptors (OTR) are particularly present in the sensory brain regions mostly used by a species to interact (olfactory in mice, auditory in songbirds, visual in macaques) [44,53]. One noteworthy exception to this pattern was the absence of OT fibers in vlPFC (area 44 and 45), neocortical substrates known to be involved in the production and perception of primate vocalizations, including marmosets [36,54,55]. A possible explanation is that OT reaches this cortical region by cerebrospinal fluid diffusion [45,56–58], though further research looking at the presence of OTR in these areas is needed. This mechanism may also explain whether the mismatch between OT fibers and OTR observed in rodents [44], and the sexually dimorphic OTR distribution [59,60], also exist in primates. Future studies are needed to address these questions, as well as the potential overlap between OT and AVP fibers, and their receptors, in the marmoset. Overall, our results suggest that OT could act at different levels of the vocal communication system, ranging from perception (auditory areas), decision making and selection of vocalizations (amygdala and area 25/32) as well as motor control (area 24), and highlight that marmosets are a keystone model in which to study the effects of OT on acoustic communication in the primate brain.

## Acknowledgments

We thank Dhananjay Bambah-Mukku for comments on this manuscript. This work is supported by grants to AL (Marie Curie-Sklodowska fellowship 101018877) and to CTM (NIH R01 DC 012087).

## Notes

### Competing Interest Statement

The authors have declared no competing interest.

